# Born to run? Quantifying the balance of prior bias and new information in prey escape decisions

**DOI:** 10.1101/297218

**Authors:** Nicholas M. Sutton, James P. O’Dwyer

## Abstract

Animal behaviors can often be challenging to model and predict, though optimality theory has improved our ability to do so. While many qualitative predictions of behavior exist, accurate quantitative models, tested by empirical data, are often lacking. This is likely due to variation in biases across individuals and variation in the way new information is gathered and used. We propose a modeling framework based on a novel interpretation of Bayes’ theorem to integrate optimization of energetic constraints with both prior biases and specific sources of new information gathered by individuals. We present methods for inferring distributions of prior biases within populations rather than assuming known priors, as is common in Bayesian approaches to modelling behavior, and for evaluating the goodness of fit of overall model descriptions. We apply this framework to predict optimal escape during predator-prey encounters, based on prior biases and variation in what information prey use. Using this approach we collected and analyzed data characterizing white-tailed deer (*Odocoileus virginianus*) escape behavior in response to human approaches. We found that distance to predator alone was not sufficient for predicting deer flight response, and have shown that the inclusion of additional information is necessary. Additionally, we compared differences in the inferred distributions of prior biases across different populations and discuss the possible role of human activity in influencing these distributions.

## Introduction

Complex behaviors such as when to search for new foraging opportunities or when to flee from predators are notoriously challenging to predict. Theoretical models based on individuals optimizing a specified quantity have made progress towards generating such predictions (Krebs et al., 1983; Parker and Smith, 1990), with optimal foraging and escape theories (Cooper Jr. and Frederick, 2007; Mangel and Clark, 1986; Pyke et al., 1977; Ydenberg and Dill, 1986) addressing the types of behavior mentioned above. What food items to consume and when to flee from predators are important problems that carry significant fitness consequences (Godin and Smith, 1988; Lind and Cresswell, 2005; Morris and Davidson, 2000; Persons et al., 2002; Thomas et al., 2001), and under optimality theory individuals should seek to optimize these behaviors with respect to fitness (Parker and Smith, 1990; Sibly and McFarland, 1976). While useful in forming qualitative predictions of behavior (Cooper Jr, 2015; Krebs et al., 1983), observed animal behavior can deviate from quantitative predictions (Krebs, 1980; Krebs et al., 1983; Pierce and Ollason, 1987; Pyke, 1984; Sayers et al., 2010), with high intraspecific behavioral variation commonly cited as an explanation for these deviations (Garamszegi and Møller, 2017; Hughes and Burrows, 1991; Pyke, 1984).

We consider two potential causes for such intraspecific variation. First, individuals may bias decision-making due to personality effects (Bergvall et al., 2011; Jones and Godin, 2010; Kurvers et al., 2009; Sih et al., 2004), the effects of prior experience, such as habituation or sensitization (Frost et al., 2007; R., 1970; Stankowich and Reimers, 2015), and internal state. Second, individuals may be limited in their capacity to acquire and utilize information, leading to decisions based on a subset of the total information available (Dall et al., 2005; Giraldeau, 1999; Koops and Abrahams, 1998). Both individual biases and variation in information use can lead to deviations from predicted optimal behaviors (Halsey et al., 2003; Hughes and Burrows, 1991; Kie, 1999; Valone and Giraldeau, 1993; Ward, 1992; Wolfe, 2013). However, due to the long-held assumption of ‘perfect information’ in behavioral ecology (the assumption that animals are cognizant of and make use of all information available to optimize decision making), models of behavior have typically ignored these sources of variation. Given that consideration of individual perception and variation in information availability can significantly alter predictions (Brown et al., 1999; Gray and Kennedy, 1994), it is important to consider both prior bias and information use when developing models.

We propose a novel interpretation of Bayes’ theorem as applied to behavior to address these two sources of variation while bypassing an assumption of perfect information. Specifically, we consider a bayesian framework in the context of antipredator behavior. There exists a large body of literature on antipredator decision-making, and in particular the decision to flee from approaching predators (Blumstein, 2003; Broom and Ruxton, 2005; Cooper and Blumstein, 2014; Cooper Jr., 2010; Cooper Jr. and Frederick, 2007; Guay et al., 2013; Møiler, 2008; Stankowich, 2008; Stankowich and Coss, 2007, 2006; Stankowich and Reimers, 2015; Ydenberg and Dill, 1986), yet few quantitative models have been developed to predict this behavior. Optimal escape theory seeks to make predictions of antipredator behaviors, such as flight-initiation distance or refuge hiding time, by comparing the fitness costs of antipredator behaviors to the risk of predation. Existing escape models (Cooper Jr. and Frederick, 2007; Ydenberg and Dill, 1986) are currently unable to quantify the effects of prior information in a meaningful way, and offer no way to identify what environmental information prey are using to make decisions (outside of testing basic qualitative hypotheses (Valone and Giraldeau, 1993)). Tests of existing models have led to some qualitative predictions (Cooper Jr., 2010; Cooper Jr, 2015), such as group A should flee earlier than group B, but no accurate quantitative predictions. We seek to provide the first case of successful quantitative predictions for optimal escape by leveraging a Bayesian inference framework to account for prior biases and variation in what information prey use.

Traditionally, Bayes’ theorem has been applied to the study of behavior in the context of Bayesian decision theory, which assumes a ‘known’ prior distribution, as opposed to Bayesian statistical inference (Dall et al., 2005; McNamara et al., 2006). In this context individuals are thought to draw from known prior distributions regarding a decision, and then update via information. For example, in foraging theory a forager faced with a patch of unknown quality may draw from the ‘known’ distribution of all patch qualities in the area to establish a prior estimate of quality, which may then be updated as foraging in the patch commences. Similarly, prey faced with a potential threat may draw from a ‘known’ distribution of risk associated with predators in the area (a sort of background predation risk) to establish a prior assessment of risk for the new threat, and then update as the encounter continues. We argue, however, that such an assumption is not necessary in an optimal decision framework, nor is it likely to be upheld in nature. Rather than consider each individual to be sampling from a known prior distribution of risk/reward, we instead think of individuals as having one prior estimation of risk/reward, based on past experience and genetic bias, that is updated after each decision making process. We then infer the distribution of these individual priors over the whole population, leading to population level descriptions of behavior that are usually impossible without extensive resampling of many known individuals over long periods of time. Following from this Bayesian approach, we establish a framework in which the conditional probabilities for updating priors may represent any number of possible decision mechanisms for interpreting different types of information during decision making.

Our modelling approach follows a general optimality and Bayesian framework (Cooper Jr. and Frederick, 2007; Mangel and Clark, 1986; McNamara et al., 2006), with some key differences as noted above, in which prey weigh the energetic costs of escape against the risk of predation. A potential attack is initiated when prey notice an approaching predator. Upon noticing an approaching predator, prey must decide whether to run, expending a known amount of energy, or remain and wait to see if the predator is going to attack, risking death but potentially avoiding unnecessary flight. In our framework, prey may update prior assessments of risk with information on any number of risk factors, such as the speed of a predator or the directness of its approach, by delaying flight in order to reduce uncertainty. Certainty that a predator will not attack will allow prey to avoid unnecessary flight, with the trade-off being that remaining to gather information may lead to an increased risk of predation. Rather than rely on assumptions of known prior distributions as is typical in a Bayesian decision theory approach to modeling behavior, we infer distributions of individual prior risk assessment at the population level. An individual’s prior is then updated based on any number of different types of information from within the encounter. The modularity afforded by generalizing to any number and type of information allows for assessment, via model comparison, of what information prey use when making decisions.

Using this approach we analyzed the escape decisions of white-tailed deer (*Odocoileus virginianus*) in East-Central Illinois. Deer are a common study species in optimal escape and flight-initiation distance (FID) studies (Stankowich, 2008; Stankowich and Coss, 2007, 2006), and readily display both vigilance and escape behaviors when approached by humans. This allows for close study of the effects of human-wildlife interactions on behavior and provides insight into how wildlife are likely to respond to actual predators. Comparing model predictions with observed escape decisions from many deer-human encounters, we find that our inferred prior distribution offers a good fit to data, passing a rigorous goodness of fit test, and enables population level behavioral comparisons. We also show that the inclusion of additional types of information can significantly improve model success without adding additional free parameters (i.e. parameters that cannot be fixed independently of the data and thus must be fitted using the data), with a two risk factor model for predicting deer flight behavior outperforming a single risk factor model.

## 1 Modeling Framework

We begin with a simple inequality comparing the costs of both possible decisions, and then expand via application of Bayes’ theorem to allow for information updating by prey during encounters. While applications of Bayesian decision theory to animal behavior in the past assume known prior distributions (McNamara et al., 2006), in wild populations tracking the effects of personality, learning, internal state, and more is not feasible, and prior distributions must therefore be inferred rather than measured directly. By utilizing a Bayesian inference approach we infer the distribution of priors in our system based on observed behavioral responses. We consider constant forms for the conditional probabilities, interpreted here as decision mechanisms for interpreting acquired information from the environment (Ydenberg, 1998), whose structure and function has been selected for over time. Using this approach we are able to compare prior distributions between populations, and can isolate the identity and form of decision mechanisms necessary for predictions of behavior.

### 1.1 Optimal escape model

We consider the following optimal escape model, where prey compare the energetic costs *β* of fleeing from a predator to the probability *P*(*A|X*) of losing energy *E:*

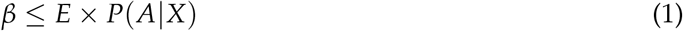

Here, *E* × *P*(*A*|*X*) is the perceived energetic cost of remaining in an encounter, represented by the product of *E,* the perceived cost of remaining in the encounter if there is an attack, weighted by *P*(*A*|*X*), the probability of attack as perceived by prey, given some information *X*. The cost to flee *β* is the metabolic costs associated with fleeing from an encounter. We consider the perceived energetic cost of remaining in an encounter if there is an attack *E* to be the daily energy budget of prey (as loss of day’s energy places individuals into a deficit, or starving, state). While true costs of remaining are much higher than daily energy budget, it is likely that *E* is based not only on some purely physiological constraints, but also on preys’ perception of how much energy they can afford to lose from any given encounter. However, we note that, qualitatively, results are insensitive to the true value of *E* over a biologically reasonable range (appendix G). When the cost of flight is less than or equal to the cost of remaining, prey should flee from an encounter. The distance between predator and prey when the cost of remaining becomes greater than the cost of flight is referred to as the optimal flight-initiation distance (FID). By considering the information *X* prey use to assess the risk of attack *P*(*A|X*) during an encounter, we can make predictions of prey FIDs.

### 1.2 Perceived probability of attack

Prey assessment of risk is the driver behind flight behavior, and the form of *P*(*A|X*) must account for what information prey use and how they might bias decisions based on prior experience. We consider prey’s estimation of risk given information *X* on the encounter, such that *P*(*A|X*) is a function of risk factors *X* that update a prior assessment of risk *P*(*A*), which we denote by *α*. Here *α* is representative of behavioral biases due to personality, habituation, and/or other effects from before the current encounter, and may be informed via observation of conspecifics (Pérez-Escudero and De Polavieja, 2011). Without identifying individuals, these effects cannot be identified specifically. Risk factors X may be related to distance to predator, or any other quantitative variable estimated by prey from observation during an encounter. *P*(*A|X*) for one risk factor is defined as follows by simply rewriting Bayes’ Theorem:

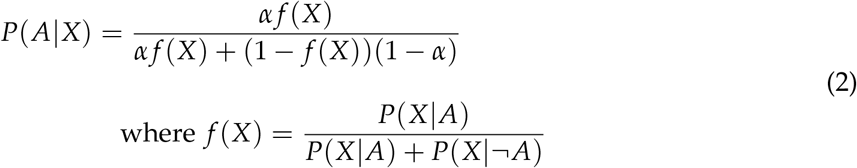

The prior *α* represents variation among individuals, while *f*(*X*) we assume to be the same for all individuals. This assumption is in keeping with the concept of decision mechanisms (Ydenberg, 1998), where *f*(*X*) represents a mechanism for interpreting acquired information, whose structure and function has been selected for over time. The probability *P*(*A|X*) can then easily be expanded to include any number of risk factors (appendix B). This approach to formulating *P*(*A|X*) provides a way for connecting decision-making with potential decision mechanisms, and is distinct from typical interpretations of Bayes’ theorem in animal behavior in two important ways. First, rather than thinking of each individual as having access to a known distribution of prior risks *α* to sample from, we think of each individual as having one singular prior, representative of the sum of past experiences, personality, and other information from before an encounter. The distribution of individual priors we then infer at the population level. Second is that we are able to generalize to any number of risk factors, and via likelihood ratio tests of models based on different risk factors and decision mechanisms we can identify what information is necessary for predictions of escape behavior versus what information is likely not utilized by prey when making decisions.

### 1.3 Inferring prior behavioral bias

The Bayesian form of *P*(*A|X*) introduces the unknown prior *α* into our model. Though the prior is unknown for any given individual, the distribution of priors in a population can be inferred from observed flight-initiation distances. We assume a beta distribution for *α*, as *α* must be between 0 and 1 and the beta distribution offers flexibility in the shape of distributions over this range. Therefore *α* is beta distributed, with shape parameters *p* and *q*, as follows:

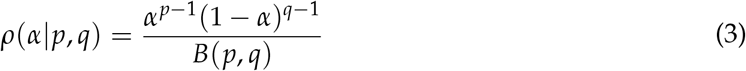

where *B*(*p,q*) is the beta function 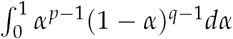. We then infer the distribution parameters *p* and *q* via maximum likelihood estimation using data from observed encounters (appendix C). Using a distribution of *α* naturally leads to predictions of prey not fleeing from certain encounters, and we therefore include both flight and non-flight observations in our likelihood calculation. By including both flight and non-flight data we are able to capture the effects of extremely bold or habituated prey that are less likely to flee. In this way the prior behavioral biases in a population, such as habituation or personality effects, can be characterized based on observed behavioral decisions. Directly measuring these unknown prior distributions, which include many different effects, is not feasible in wildlife populations and such effects must be inferred instead. To date, the ability to capture these effects to this degree has been absent from optimal decision making, escape models, and FID analyses.

### 1.4 Model comparison and goodness of fit

Using our inferred distributions of *α* we conduct an exact test on the goodness of fit of our model. We begin by generating a large number of predictions of encounter outcomes based on known encounter information such as approacher speed and directness, with prior *α* drawn from our inferred distribution. We then generate a distribution of likelihoods from the generated data sets. By comparing the likelihood of our observed FIDs and non-flight data from the field to the distribution of likelihoods generated using our inferred *α* distribution we determine whether our model is a good fit for describing the observed flight responses. This test enables us to assess how likely any given model is to generate a data set similar to what we have observed, providing us with a measure of accuracy regarding model predictions. After identifying potential models that offer good fits to our data, we then compare models via likelihood ratio tests to select the best model, allowing us to determine whether including or excluding certain types of information will yield better predictions.

## 2 Methods

We tested our modeling framework in the context of white-tailed deer escape decisions during encounters with approaching humans. The energetic parameters in our model were set using deer metabolic data from the literature, and observed deer escape behavior was compared to predictions from models considering either distance to predator alone, or distance and directness of approach together as risk factors.

### 2.1 Sampling

Deer flight-initiation distances (FIDs) were collected following (Sutton and Heske, 2017) but with varying angles of approach (see appendix H for a detailed description of approach method). From March through August 2016 deer at Kickapoo State Park (Vermilion County, IL, USA, 40.1167° N, 87.7544° W) were approached randomly at 0, 10, 20, 30, 40, and 60 degrees and FIDs were recorded using a Nikon Prostaff 3i laser rangefinder (Nikon, Inc., Melville, NY). The observer, N.M.S, wore the same attire and maintained a constant velocity of 1 meter/s during all approaches so as to reduce unwanted variation in approaches. Data was censored if encounters were interrupted at any point by other park visitors. Because individuals could not be identified, some individuals may have been resampled (we estimate the probability of resampling any one individual to be approximately 0.012 given park size and estimates of deer population in the area). However, past studies have shown that a moderate degree of pseudoreplication will not significantly alter the results of most FID analyses (Runyan and Blumstein, 2004). Additionally, the effects of individual identity are a part of prior *α*, which are included via maximum likelihood inference rather than measured directly.

### 2.2 Model parametrization

White-tailed deer energetic parameters daily energy budget (Nixon et al., 1991; Thompson et al., 1973), flight cost (Mautz and Fair, 1980; Nixon et al., 1991; Sweeney et al., 1971), and deer top speed (Garland and Janis, 1993) were estimated from the literature. Flight cost was estimated to be 135 kcal, daily energy budget was estimated to be 5500 kcal, and deer top speed was estimated to be 18 m/s. Flight cost was calculated considering species-specific metabolic costs of locomotion and average length of chases when confronted with predators (appendix D). These values are subject to variability within any wildlife population and therefore represent a best guess at the average values for the deer in our study.

### 2.3 Describing encounters

We consider deer-human encounters in a polar coordinate system, similar to past studies of escape behavior (Broom and Ruxton, 2005), with deer immobile at the origin, as shown in Figure 1. An encounter begins when deer detect an approaching human at alert distance AD. The approacher approached deer directly until they exhibited alert behavior, at which point the approacher assumed an approach angle at AD. The human continued approaching the deer following a linear path angled *θ* degrees away from the deer and with constant velocity *V* = 1 m/s. At any time t the distance between human and deer *r*(*t*) and the component of human velocity towards the deer *dr/dt* may be calculated. The distance *r*(*t*) when the deer flees is the flight-initiation distance *FID*.

**Figure 1:**
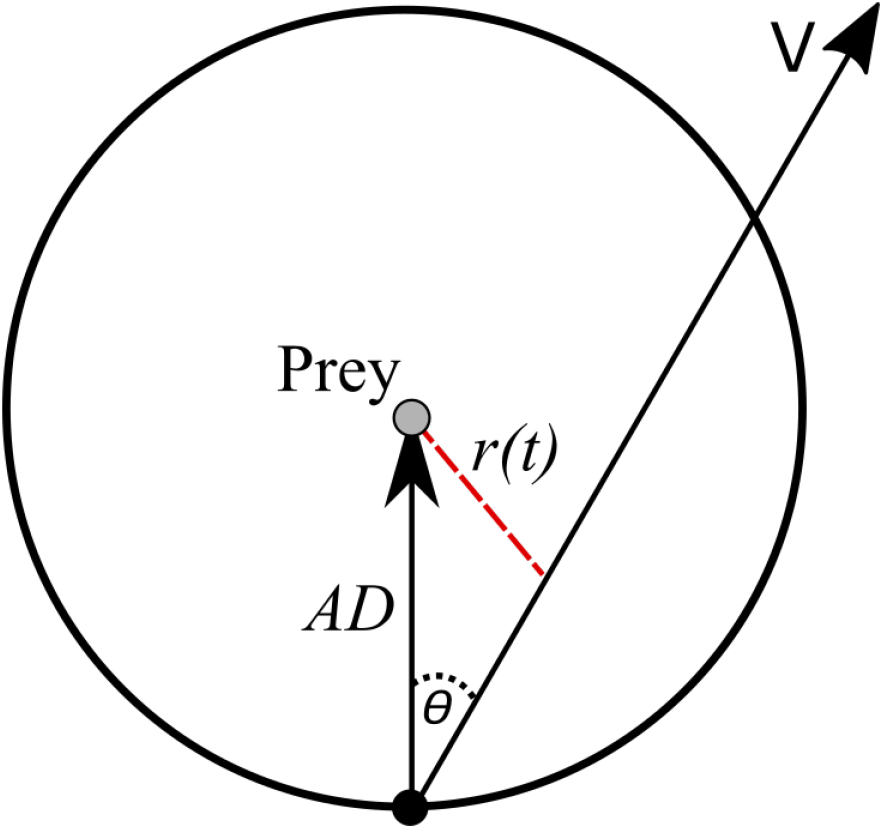
Top-down, graphical representation of encounters between humans and deer. Here, the circle represents the encounter space and is defined by the distance when the deer first spots an approaching human, also know as the alert distance *AD*. Beginning on the edge of the encounter space *AD* meters from the deer at the origin (labeled prey in this figure) a human would then approach the deer along some path indicated by the arrow labeled *V*, at some angle *θ* and with constant velocity *V*. Distance between observer and deer *r*(*t*) can be calculated at any time t along the path. The derivative of *r*(*t*) here is the component of *V* in the direction of the prey.

### 2.4 Risk factors

In the absence of bias, or equivalently when *α* = 0.5, we find that *f*(*X*) = *P*(*A|X*). Given that *P*(*A|X*) should interpolate between zero and one in some limits of the value of *X*, we develop an ansatz for the form of *f*(*X*) in the absence of bias. We consider two functions or decision mechanisms, *f*_1_(*X*) and *f*_2_(*X*) for two risk factors; distance to predator, the main factor considered in existing models, and directness of predator approach, which empirical evidence suggests is important in mediating escape decisions (Guay et al., 2013; Stankowich and Coss, 2007, 2006), respectively. For *X* equal to distance to predator *r*, we estimate perceived risk of attack *f*_1_(*r*) such that risk increases as *r* increases linearly relative to the initial distance *AD* at which the predator is detected:

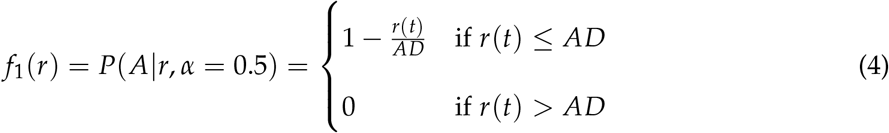

By considering distance to predator relative to AD, rather than absolute distance only, we propose a decision mechanism that accounts for prey analyzing how much a predator has moved towards them since initial detection. In contrast, if prey only considered current distance to predators when assessing risk, then it would suggest no consideration for the information gathered from predator movement since initial detection. Additionally, a dependence on *AD* in the distance decision mechanism effectively creates a reasonable lower bound on risk. The alert distance sets the stage for the encounter, such that if a predator moves further away so that *r* becomes greater than *AD,* the predator can be thought of as leaving the encounter and risk becomes zero. For *X* equal to directness of approach we consider *f*_2_(*v*_*r*_) as increasing linearly with increasing velocity in the direction of the prey *v*_*r*_, relative to the maximum velocity of the prey *v*_*max*_. Increased angles of approach result in *v*_*r*_ approaching zero, when observer position is tangent to deer at origin, at a greater rate:

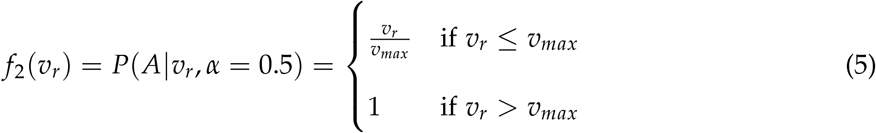

Our rationale for defining *v*_*max*_ as we have is that the perceived risk of predation should reach a probability of one as soon as prey can no longer outrun an approaching predator. While more general forms for these decision mechanisms exist, we chose a linear relationship between risk factor and risk as the simplest first step for defining these factors. We also note that adding additional decision mechanisms and risk factors, such as predator posture or directness of gaze, is relatively easy and adds no free parameters to the model. This is because each decision mechanism is merely an interpretation of data present during encounters, such as *r* and *AD*, or else bioenergetic parameters that can be estimated from the literature or measured directly, such as *v*_*max*_.

### 2.5 Analyses

We performed an exact test (Etienne, 2007) to test the goodness of fit of our models and inferred prior distributions, and Vuong’s closeness test, a likelihood ratio test for non-nested models, to compare and select the better of our one and two risk factor models (appendix F). Our goodness of fit test effectively checks for overfitting, while Vuong’s closeness test takes into account the effect of additional free parameters, if we had any, when comparing the likelihoods of different models. All analyses were performed at the 0.05 significance level. Analyses were performed in R version 3.3.0 (R Core Team, 2016). Data and code are available on the O’Dwyer Lab GitHub repository (https://github.com/odwyer-lab/BornToRun).

## 3 Results

We visited Kickapoo State Recreation Area 11 times during our study and recorded ADs, FIDs, and flight/non-flight data for 40 encounters with deer over six angles of approach. Eleven deer were approached directly at zero degrees, while the remaining 29 deer were approached indirectly. Of these indirect approaches, 9 were at 10 degrees, 9 at 20 degrees, 7 at 30 degrees, 1 at 40 degrees, and 3 at 60 degrees. From 30 degrees and above, the majority of encounters resulted in non-flights.

### 3.1 Model goodness of fit and risk factor comparison

Our two risk factor model accurately described observed flight behaviors, with FID distributions and proportions of non-flight predictions tracking variation across angles of approach and between individuals (Figure 2). The two risk factor model based on distance to predator and angle of predator approach passed an exact test on the model log likelihoods (*p* = 0.6; Figure 3) when predicting FIDs and proportion of non-flights. A one risk factor model, with angle of approach as a risk factor removed, also passed the exact test (*p* = 0.21; Figure 3). When comparing the two models, Vuong’s closeness test (a likelihood ratio test) indicated that our two risk factor model was significantly closer to the true data generating process than the one risk factor model (*z* = 2.34, *p* = 0.01). These results suggest that the inclusion of additional risk factors, such as directness of approach, can lead to models of flight behavior that significantly outperform models based solely on distance to predator as a risk factor. The inclusion of additional risk factors is therefore vital, especially when approach paths are varied.

**Figure 2:**
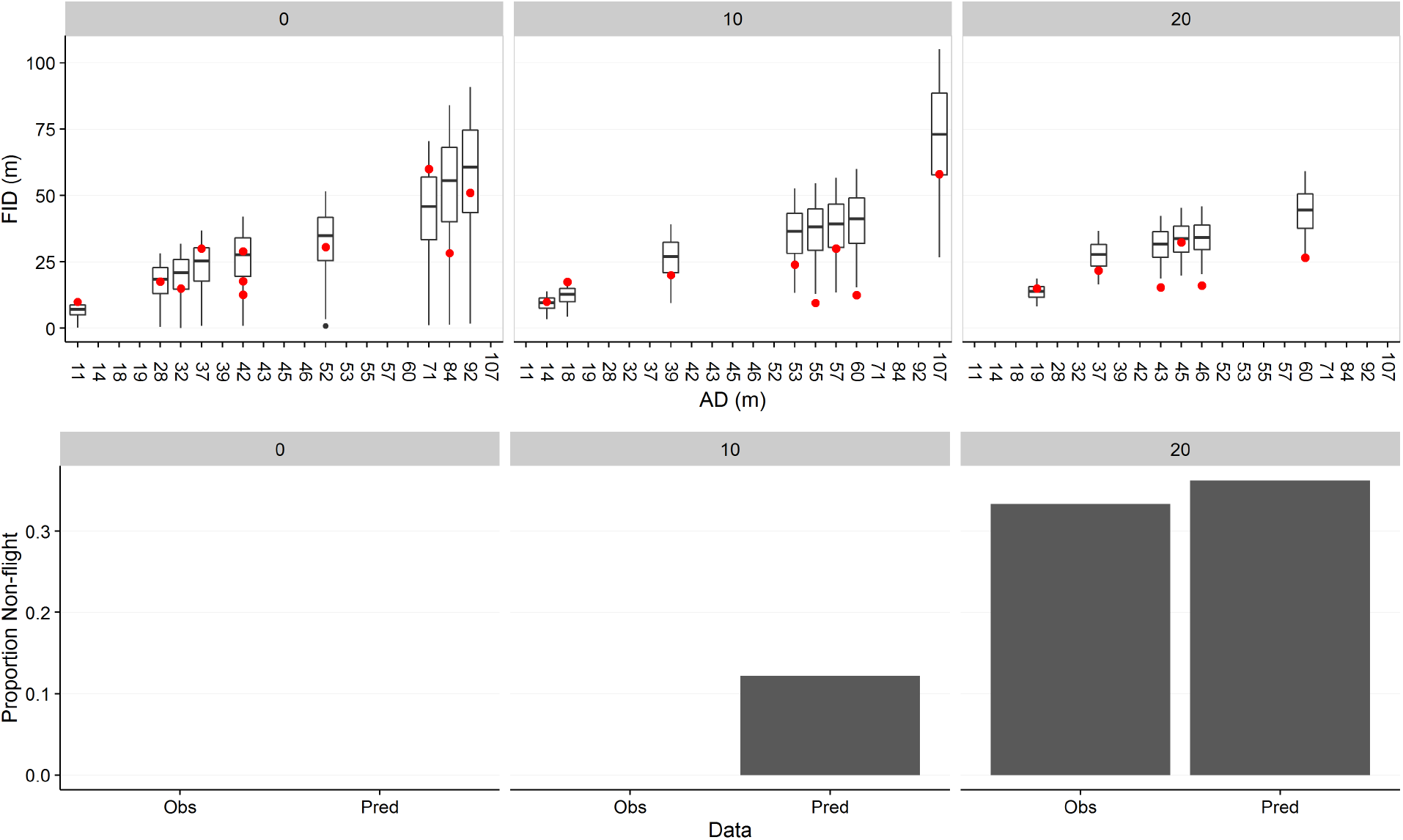
**Top**: The top three panels show observed FIDs (red points) and model predicted FIDs via boxplots for 0, 10, and 20 degree approaches, with alert distance AD in meters on the x-axis. Observed FIDs fall within predicted FID distributions, suggesting that the model is doing well at predicting FIDs while accounting for the significant variability in escape behavior. **Bottom**: The bottom three panels show observed versus predicted proportion of encounters resulting in non-flight for 0, 10, and 20 degree approaches. Model successfully predicts proportion of non-flights across angles, with the largest discrepancy at 10 degrees where no non-flight is observed but non-flight is predicted approximately 10% of the time (i.e. for 90% of encounters at 10 degrees the model successfully predicts flight should occur). These results were produced following equation 1, using the decision mechanisms described in equations 4 and 5.

**Figure 3:**
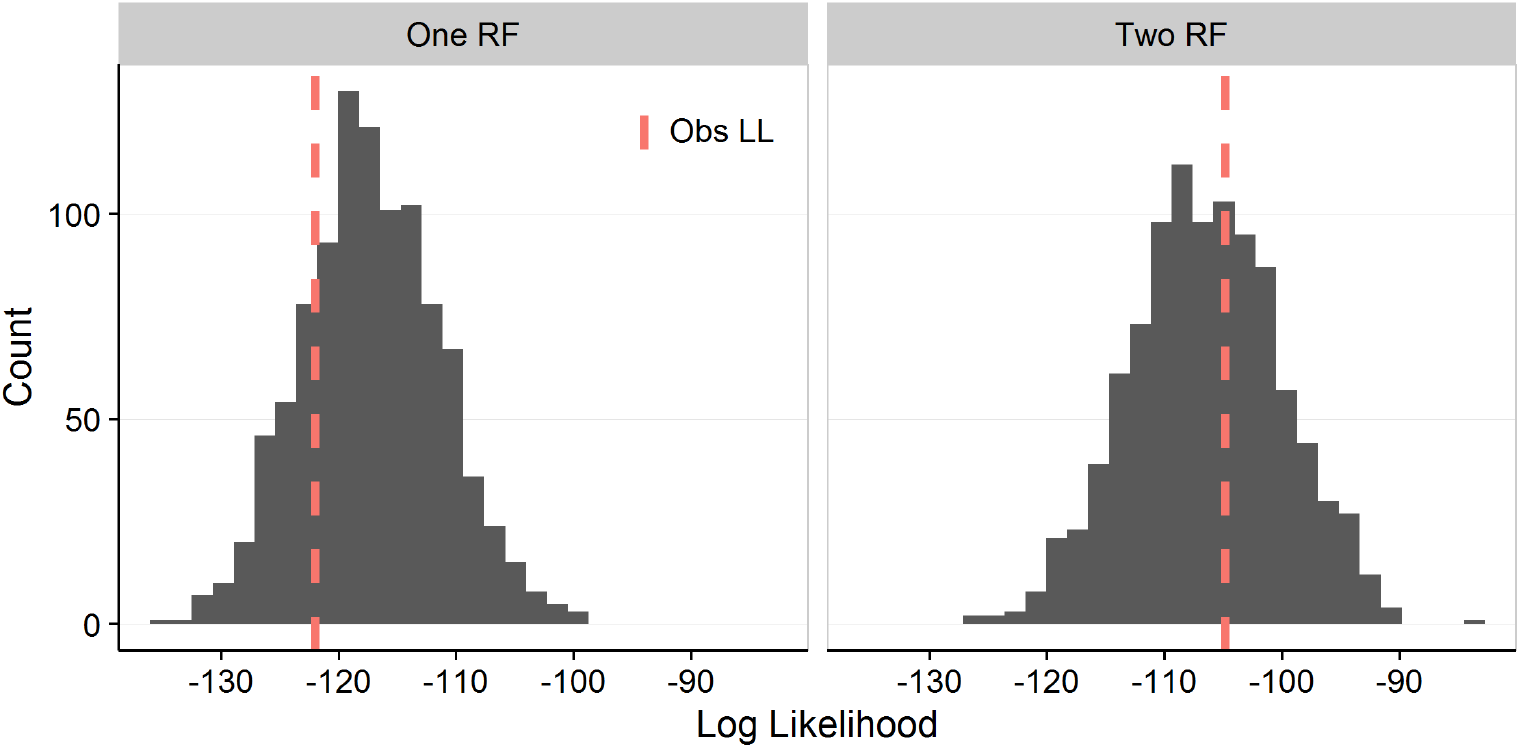
Distribution of log likelihoods for predicted data sets of flight behavior given *α*’s randomly sampled from our inferred *α* distribution. These likelihood distributions are generated to measure the accuracy of model predictions via a goodness of fit test. Red, dashed line indicates log likelihood of observed data set and is used to determine how likely observed flight behavior is given what the model is capable of describing. **Left**: Observed data was in the 21^*st*^ percentile of one risk factor model LLs. **Right**: For the two risk factor model, observed data was in the 60^*th*^ percentile of the generated LL distribution. Both models pass our goodness of fit test, meaning that the typical outcome of our inferred models is consistent with the observed data.

### 3.2 Population level inference and comparison of prior distributions

The inferred α distribution for our study site, Kickapoo State Park, suggests significant variability in deer prior biases (*µ* = 0.46, *σ*^2^ = 0.05; Figure 4A). While we know that this inferred distribution, in conjunction with our implementation of information gathering, provides a good fit to observed data, the interpretation of the absolute value of ***α*** is somewhat subjective given the flexibility of our model to adjust for changes in the functional form of our risk factors. For a meaningful interpretation of our study site’s *α* distribution we compare to a second population from a different study site. In a prior study examining the effect of variation in park use on deer escape behavior, deer in our study site, Kickapoo State Park, were found to be the most habituated and/or bold of all parks studied, allowing humans closer than at other parks (Sutton and Heske, 2017). Deer in Moraine View State Recreation Area, a different site which typically receives significantly fewer human visitors per year than Kickapoo, fled earlier in encounters. Using past data from deer encounters at Moraine View we inferred an *α* distribution with less variance than Kickapoo (*µ =* 0.58, σ^2^ = 0.03; Figure 4B). By comparing the mean and variance of *α* for these two sites we can see that deer in Moraine View are shifted towards larger *α* values, indicative of sensitization and/or shy personalities and biasing decisions towards earlier flight. Deer in our study site, Kickapoo, are shifted toward intermediate or smaller *α* values, suggesting they are more habituated and/or there are more bold deer in Kickapoo than in Moraine View. In this way, we are able to link variation in human land use to population level variability in behavioral priors.

**Figure 4:**
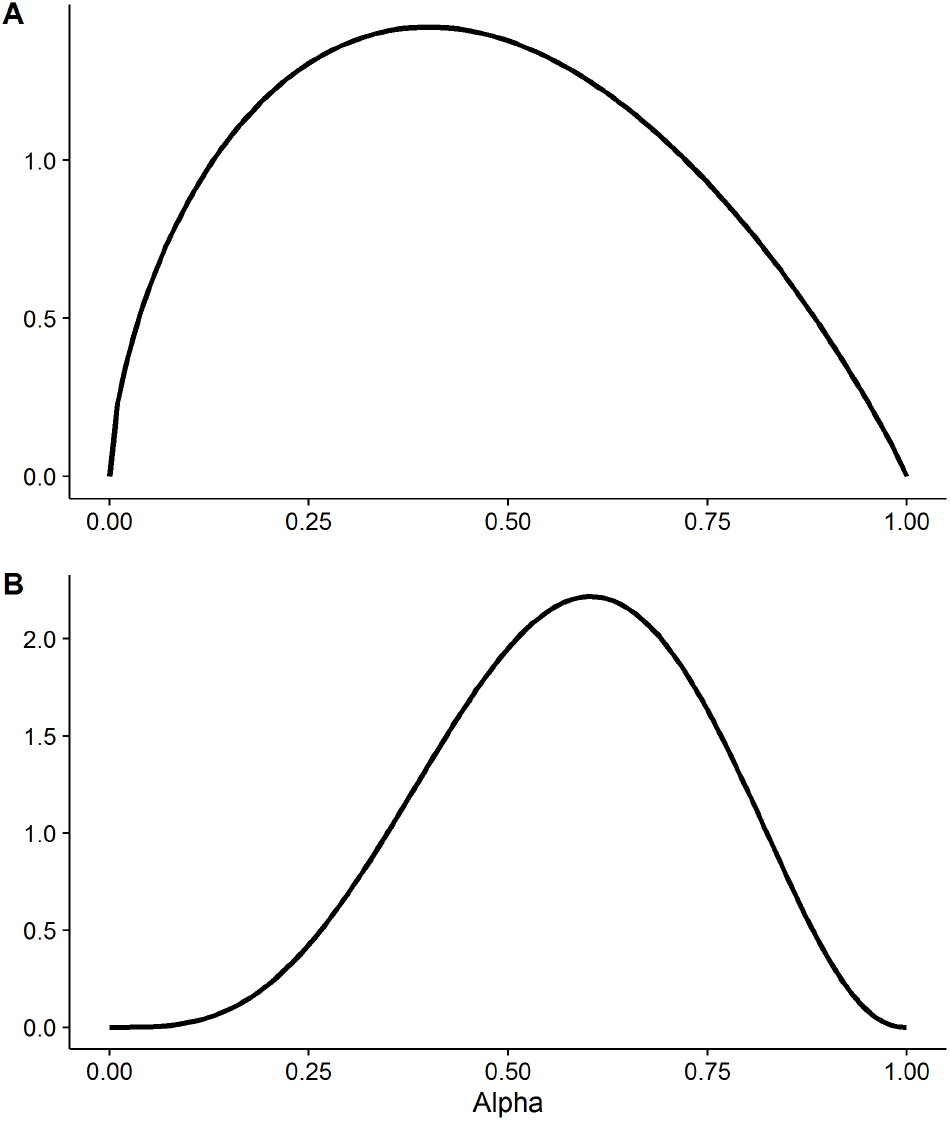
Inferred distribution of *α,* a measure of prey prior bias and indicative of habituation level and/or personality, for Kickapoo State Park (A) and Moraine View State Recreation Area (B) deer populations. Large *α*’*s* correspond with more sensitized and/or shy individuals, whereas small *α’s* correspond to more habituated and/or bold individuals. Deer in A (*µ* = 0.46 and *σ*^2^ = 0.05) are shifted towards habituation, or bold personality, relative to B (*µ* = 0.58, *σ*^2^ = 0.03). Deer in B are shifted towards sensitization, or shy personality.

### 3.3 Additional analyses

To test the sensitivity of our results to the choice of *E*, we additionally considered values of *E* ranging from a lower limit of *E* equal to cost to flee, up to an estimation of the total caloric value of a deer, around 76000 kcal (Weiner, 1973). Qualitatively, results were insensitive to this choice for *E*, and in fact a general trend of decreasing model log likelihood was observed as *E* increased, providing support for choosing a smaller value of *E* (appendix G). Previous studies have fitted a linear model to describe the relationship between alert distance (AD) and FID, and so we have also included a regression analysis for comparison, with some caveats regarding the appropriateness of such a comparison in this context. We find that such an approach ignores sources of variation in FID and does not perform as well as our modelling approach (appendix *E*).

## Discussion

Our framework includes methods for analyzing the importance of different types of predator cues and information for making escape decisions, and shows that the inclusion of additional types of information in an optimal escape model can significantly improve its predictive power without adding additional free parameters. A model using distance to predator as the only risk factor for predicting escape behavior, as is common in existing optimal escape models (Cooper Jr. and Frederick, 2007; Ydenberg and Dill, 1986), performed significantly weaker than a competing model that included angle of approach as an additional risk factor. The purpose of comparing our one and two risk-factor models is to determine whether, in encounters where both distance and angle of approach vary, prey use only one or both variables for risk assessment. From empirical studies of escape behavior we know that additional factors such as approacher speed, directness of gaze, posture, etc. can all affect escape behavior, as can demographic factors such as group size, age, and sex (Stankowich, 2008; Stankowich and Coss, 2007, 2006). The inclusion of any of these as additional risk factors might further improve model accuracy, as well as allow for predictions under more variable approach conditions. The nature of how these factors affect prey perception of risk is still poorly understood. While we assume simple linear relationships between risk factor and perceived risk of predation as a first step, more general forms of these decision mechanisms would allow for more complicated, nonlinear relationships, though at the cost of additional free parameters.

We characterized the behavioral variation within populations via maximum likelihood inference, assuming a beta distribution for prior *α*. This parameter, *α*, is representative of the cumulative effects of personality, prior experience, internal state, and any other processes involved in biasing initial assessments of predation risk. These biases are important for understanding the effects of individual behavioral variation on escape behavior, as high intraspecific behavioral variation is commonly cited as an explanation for discrepancies between observed and predicted behaviors (Hughes and Burrows, 1991; Pyke, 1984). Our inferred distribution implies moderate habituation to humans throughout our study site and/or a slight shift towards bolder personalities when compared to a previous site, consistent with known extents of human land use at both sites. Furthermore, our exact test indicates that these distributions represent a good fit to our observed data. It is possible that a different distribution may offer a better fit, especially if subsets of the population differ in behavior, so it may be worth considering alternative distributions. Our approach to inferring population level behavioral biases affords us the ability to rapidly describe population level behavioral characteristics, using a small amount of data, that would be otherwise impossible without extensive tracking of individual behaviors through time. However, a downside is that the mechanisms underlying this variation cannot be parsed out, as identifying whether variation is due primarily to personality versus habituation or other effects would require repeated measures of individual behavior over time.

Taken together, the results of our inference of behavioral priors and characterization of risk factors reveal several important advances in modeling decision making behaviors. Our results demonstrate that a framework with an unknown but inferrable distribution of priors in a population, in conjunction with information updating, not only works but offers a good description of observed behavioral responses. Following a Bayesian inference approach provides both a method for inferring unknown priors and, via a likelihood ratio test, a quantitative way for testing different hypotheses of what decision mechanisms drive information updating in our system. This approach to describing decision-making circumvents the assumption of perfect information found in Bayesian frameworks where priors and the information used for updating are assumed to be known. Furthermore, our framework provides a general method by which empirical evidence and natural history expertise may be incorporated into models via any number of risk factors *X* and associated decision mechanisms *f*_*i*_(*X*) that may be worth investigating, simply by utilizing animal behaviors observed in the field.

The ability to capture the effects of behavioral variation and to easily incorporate additional types of information into models is key to improving the accuracy of quantitative models of behavior, and towards this end our approach offers significant progress. Our framework provides support for Bayesian inference methods in animal decision-making, in contrast to the tendency of assuming prior distributions, and sets the stage for broader behavioral analyses and comparisons. The good fit offered by our behavioral prior inference shows promise in comparing distributions of habituation levels, variation in internal state, differences in personality, and more across wildlife populations, with potential for analyses of these effects on population level patterns and dynamics. Rather than assume what information is used in updating priors, our risk factor/decision mechanism approach aids in quantifying information use and opens the door for further questions regarding choice of information when making decisions, variation in information use across populations, and even co-evolutionary strategies between prey information use and decision making and predator approach strategy.

## Acknowledgments

This work was supported by the Cooperative State Research, Education, and Extension Service, US Department of Agriculture, under project number ILLU 875-952. J.O.D. acknowledges the Simons Foundation Grant #376199 and McDonnell Foundation Grant #220020439.

## Online Appendix

### A Identifying potential risk factors

We used analysis of variance (ANOVA) and post-hoc Welch’s unequal variances *t*-test to test for the effects of alert distance (AD) and angle of approach *θ* on deer FIDs to reaffirm the effects of these variables and to ensure they are worth modeling.

Deer FIDs varied across angles of approach *θ* (*F*_1,38_ = 8.83, *p* = 0.005; and see Figure *A*1 for visualization of FID versus Angle of approach). FIDs were significantly smaller for indirect approaches versus direct approaches (*t*_16_ = 2.2843, *p* = 0.036), consistent with past studies. Furthermore, the proportion of encounters resulting in non-flight differed across angles (*F*_1,38_ = 33.45, *p* < 0.0001) and was significantly higher for 30 degree approaches and above than for approaches less than 30 degrees (*t*_20_ = 2.7385, *p* = 0.01).

**Figure A1:**
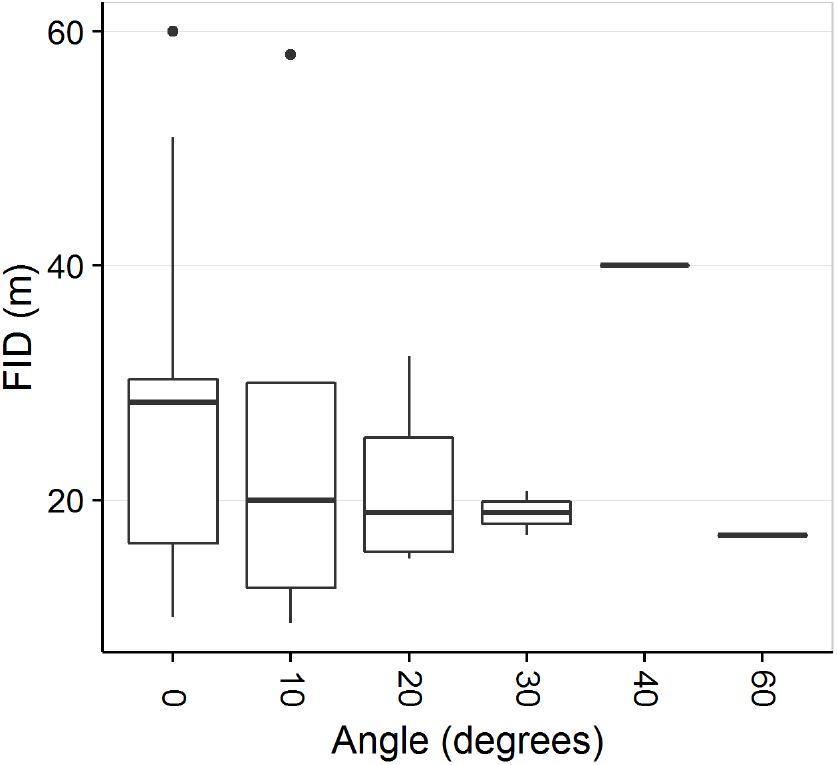
Boxplots of FIDs across the angles of approach sampled. The box plots summarize the distribution of flight initiation distances as a function of approach angle. At 40 and 60 degree angles, all but one approach each resulted in non-flight (i.e. the boxplots shown for these two angles are based on a single, atypical, encounter). The typical FID reduces with increased angle (this is consistent with velocity as a risk factor, given the dependence of radial velocity on angle of approach).

### B Perceived probability of attack

We begin by considering prey’s perceived probability of being attacked given one risk factor, or one type of information *X*, following Bayes’ rule. Here, *α* acts as the prior assessment of risk, and is representative of prior biases such as those due to habituation/sensitization, personality type, and/or other processes that may bias behavior prior to information gathering:

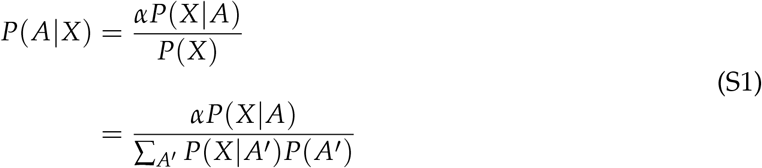

As the event *A* is binary, there is either an attack or not an attack, this sum expands to:

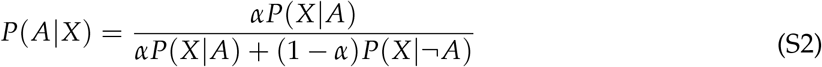

We now divide through by *P*(*X|A*) + *P(X|¬A*) in order to express *P*(*X|A*) and *P*(*X|¬A*) in terms of one function of information *X*, *f* (*X*):

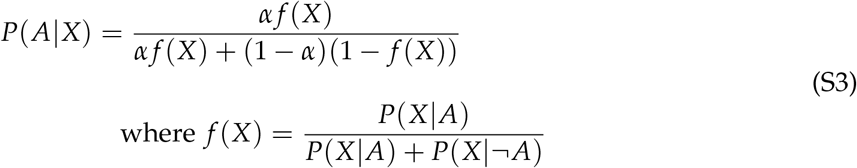

One way to interpret the meaning of *f*(*X*) is to note that when prior probability *α* = 0.5, *P*(*A|X*) = *f*(*X*):

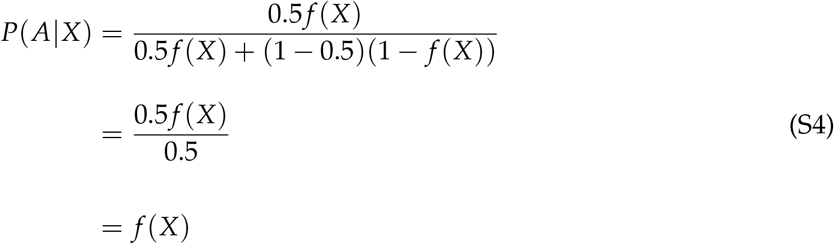

#### B.1 Multiple risk factors

For expanding to multiple risk factors we first consider adding one additional risk factor, for a total of two, and assuming that *X*_1_ and *X*_2_ are independent conditioned on *A*:

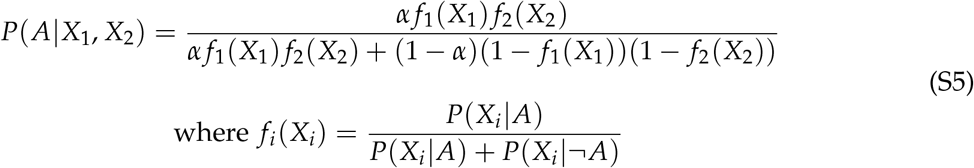

An assumption of conditional independence for our risk factors is not unreasonable if *f*_1_(*X*_1_) and *f*_2_(*X*_2_) are viewed as separate decision mechanisms (Ydenberg, 1998) for analyzing the two risk factors, and that these mechanisms may have evolved independently. We can then expand for any *n* number of risk factors:

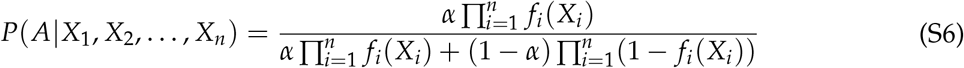

When *α* = 0.5, the probability of attack given all risk factors of interest is equal to the product of each independent assessment of risk *f*_*i*_(*X*_*i*_):

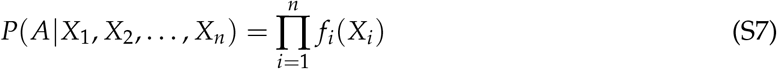

### C Maximum likelihood estimation

We begin by considering the likelihood of observing our flight and non-flight data for a general probability distribution of *α*, indicated by distribution parameter *θ*:

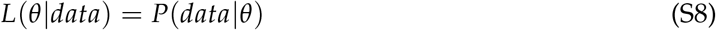

We then assume a beta distribution for *α*, as *α* must be between 0 and 1 and the beta distribution offers flexibility in the shape of distributions over this range. Our likelihood equation is updated to incorporate shape parameters *p* and *q*:

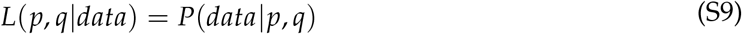

We now use maximum likelihood estimation to infer shape parameters *p* and *q* for a beta distributed prior *α* given observed flight initiation distances, denoted here simply by *F*. Following is our log likelihood equation (*n f* refers to non-flight data and lower-case *f* refers to flight data. Additionally, please note that we have dropped the subscript i from *α*_*T*_, *α*_*L*_, and *α*_*U*_ to avoid clutter). Our log likelihood equation has an additional term for summing over non-flight data via the regularized incomplete beta function. This is because *α*(*F*) cannot be calculated directly for non-flights, as a range of *α* values can result in non-flight for these instances:

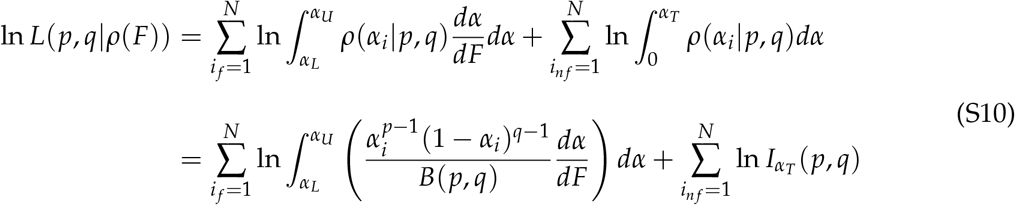

where *α*_*T*_ is a threshold value of alpha, below which flight never occurs for a given encounter, and is representative of the closest point *F*_*T*_ to prey along a given path (i.e. closest an approacher could get to prey given angle of approach; given by 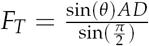), and *I*_*αT*_ (*p, q*) is the regularized incomplete beta function:

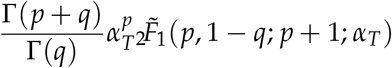

We integrate the first term in the likelihood equation over a small range to account for error in observed data and instances where *α* changes very quickly. Here *α*_L_ and *α*_U_ are the lower and upper limits of integration, defined as follows:

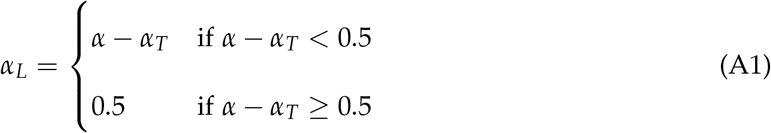

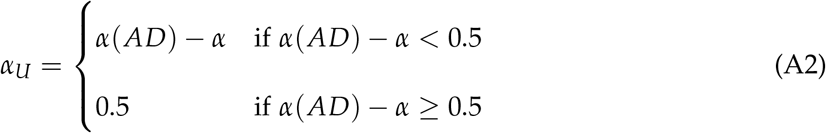

As most observations result in intermediate values of *α*(*F*) between *AD* and *F*_*T*_, the range of integration is typically one, which is a close approximation to the likelihood when no effect of error is assumed (i.e. when the likelihood is taken at a single point rather than integrating over some small range; in fact, the likelihood is relatively insensitive to the inclusion of error). In instances where flight occurs very near AD or *F*_*T*_, the range of integration is smaller to avoid integrating over values outside the bounds of the encounter (where *AD* and *F*_*T*_ are the upper and lower bounds of an encounter). Shape parameters *p* and *q* are then estimated via numerical likelihood optimization using the BFGS algorithm in R Studio via the optimx package (Nash, 2014; Nash and Varadhan, 2011).

### D Flight cost *β*

Cost to flee *β* is estimated from (Mautz and Fair, 1980):

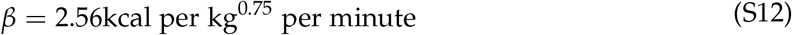

and was calculated using estimates of kg (Nixon et al., 1991; Thompson et al., 1973), top speed (Garland and Janis, 1993), and average distance of chases (Sweeney et al., 1971). Over the course of encounters lasting less than a minute, missed opportunities costs in the form of foraging during encounters were estimated to be insignificant, and therefore were dropped from our *β* calculation.

### E Comparison with linear modelling approach

Previous studies of escape behavior have noted the approximately linear relationship between start or alert distance (AD) and FID (Blumstein, 2003; Cooper and Blumstein, 2014), prompting us to consider a general linear model as an alternative approach (though such linearity may be a mathematical artefact given FID is necessarily less than AD, and a linear modeling approach might therefore be inappropriate (Dumont et al., 2012)). We constructed the following linear model to predict FID given alert distance *AD,* angle of approach *θ*, and their interaction. We performed a multivariate multiple regression analysis and post-hoc multiple analysis of variance (MANOVA) on this general linear model:

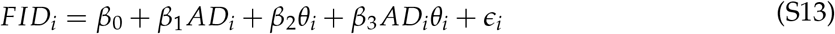

For easy comparison to the results of the multivariate multiple regression analysis, we calculated the correlation between our optimal escape model predictions and observed behavior for FID and non-flights. As our model predicts a distribution of FIDs per *AD* and *θ* we used mean FIDs for this correlation. It is important to note that a linear model for predicting FID cannot account for non-flight behavior, whereas our optimal escape model can simultaneously predict both FID and non-flight behavior. Additionally, that our new modeling framework generates a distribution of predicted FIDs per encounter allows us to capture the significant behavioral variation inherent in any wildlife population. Attempting to condense this variation into a correlation coefficient by using mean predicted FIDs effectively ignores one of the main benefits of our approach.

The general linear model with AD and angle of approach *θ* as predictors of FID explained a significant amount of variation in FID (*R*^2^ = 0.52, *p* < 0.0001), suggested no effect of angle (*p* = 0.52), and could not provide an analysis for non-flight behavior. For comparison, regressing observed FIDs and non-flight behavior on the mean of our model FID predictions and proportion of non-flights explains more variation while also accounting for non-flights (*R*^2^ = 0.84, *p* < 0.0001). While the previously identified approximately linear dependence of FID on AD might suggest fitting a linear dependence on both FID and approach angle here, this approach fails to capture the variance in the data to the degree that our model does, and cannot account for non-flight behavior or detect the effect of angle.

### F Vuong’s closeness test

As our one and two risk factor models have the same number of free parameters, we needed an analysis for comparing non-nested models. We used Vuong’s closeness test, a likelihood ratio test capable of handling both nested and non-nested models. The test compares the difference in the maximum likelihood of the two models of interest (corrected for number of parameters in each) to the mean squared pointwise log likelihood ratios. For our two non-nested models, Z-statistic is calculated as follows:

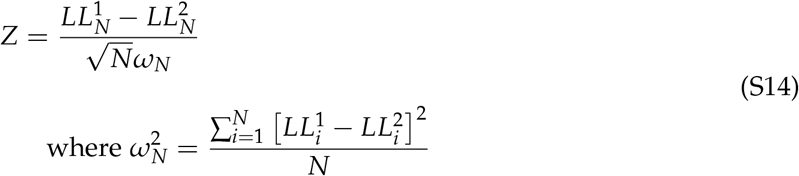

If *p* < 0.05 we fail to reject the null that both models are equally close to the true data generating process, and conclude that model one is closer. This test effectively provides a measure of significance for determining which of our two models performs better.

### G Results are qualitatively insensitive to value of E

We argue that the value of *E* in our system is equivalent to the daily energy budget of deer, as this is representative of the energy immediately available for ‘gambling’ so to speak. However, the true value of *E* is likely much larger as death by predation would lead to loss of all energy, a value much higher than the daily energy budget due to sources such as fat reserves, not to mention the energetic value of potential future offspring. Regardless of the true value of *E*, we find that our results are qualitatively insensitive to choice of *E* over a biologically reasonable range of values. If we take a value of *E* an order of magnitude larger than our daily energy budget estimate, *E* = 76287.183 (which we estimate to be the total caloric value of a 35.3kg deer (Weiner, 1973)), Vuong’s closeness test still finds the two risk factor model to be significantly better than the one risk factor model (*z* = 1.72, *p* = 0.04) and both one and two risk factor models still pass a goodness of fit test (*p* = 0.13 and *p* = 0.28, respectively). Furthermore, if we treat *E* as an additional free parameter to be fitted, we find that the maximum log likelihood of the model remains largely unchanged (and in fact the likelihood even decreases a little, though not significantly so; Figure A2). This suggests that the fit of the model is relatively insensitive to the true value of *E*. We do wish to note, however, that as *E* increases to infinity, all risk factors will effectively drop out of the model and decisions will be entirely dependent upon prior *α* only (and as such model risk factor comparison at this limit is no longer meaningful).

**Figure A2:**
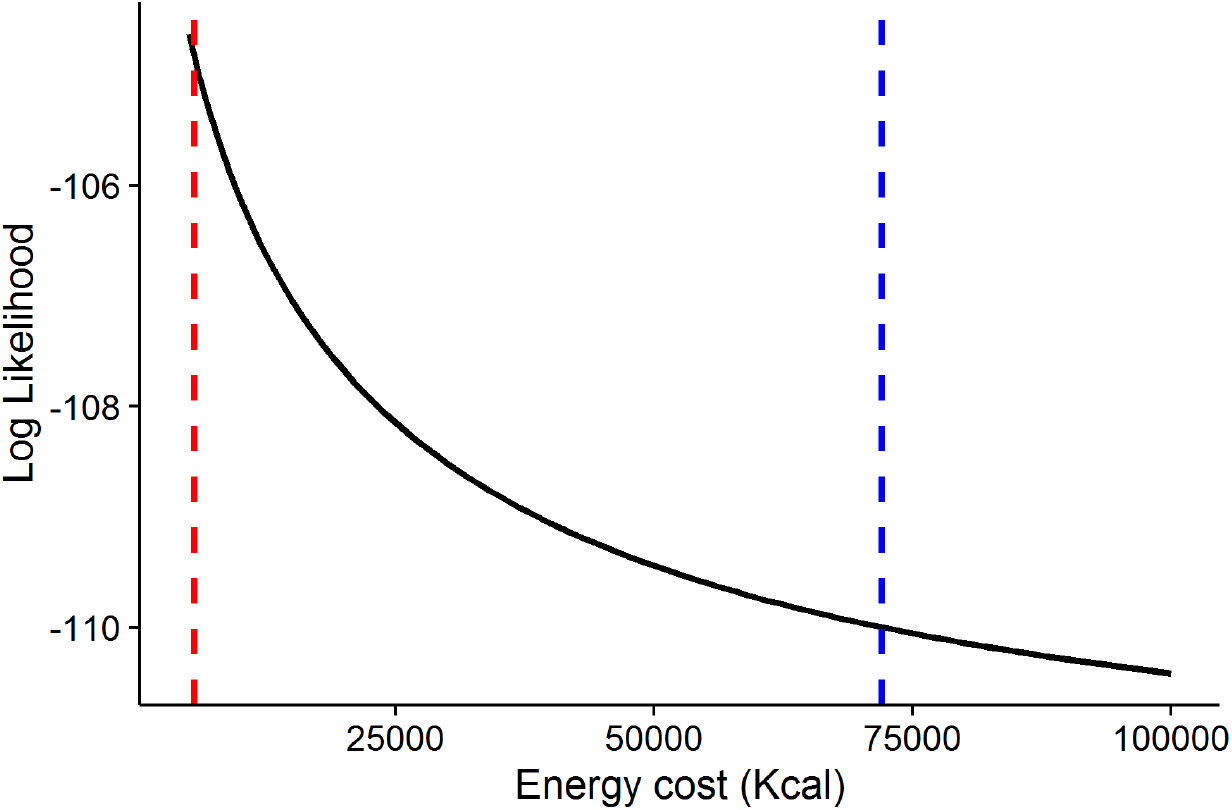
Model log likelihood as a function of cost of remaining *E*. Cost *E* was treated as a free parameter, along with beta distribution shape parameters *p* and *q*, and was fitted via maximum likelihood estimation. We can see that over a large, biologically relevant range of values for *E* (from estimate of daily energy budget, indicated by the leftmost, red-dashed vertical line, to estimate of total caloric value of a deer, the rightmost, blue-dashed line) the log likelihood remains largely unchanged, with a trend toward decreasing model likelihood as E increases.

### H Additional sampling methods

During each site visit, the approacher began searching for deer one hour before sunset and ended the survey 30 minutes after sunset to control for time of day as a factor affecting encounter rates and escape behavior. Surveys were conducted primarily in high-traffic areas of parks (e.g., roadsides, hiking and horseback trails) to ensure that any deer encountered were likely to be deer that also encountered park visitors. The approacher wore the same attire (khaki cargo pants, long-sleeved grey button-up shirt, and grey beanie) for every encounter. Once deer were located, the approacher selected a focal deer if the deer occurred in a group and positioned himself with a clear, straight-line path to the deer. The approacher used a weighted flag to mark this as the initial, starting distance (SD), then walked towards the deer at a constant speed. When the focal deer became alert (head upright and pointed in the approacher’s direction), the approacher dropped a second weighted flag to mark the alert distance (AD). At this time, the approacher quickly referenced a protractor attached to the clipboard/datasheet to assume one of six approach angles (0, 10, 20, 30, 40, or 60 degrees) and continued approaching along this trajectory. This was done to control the angle of approach starting from when deer first became alert (as starting angled from SD leads to significantly more variable angles of approach from the deer’s perspective, depending on AD). Finally, when the deer fled the approacher dropped a third weighted flag to mark the FID. The approacher then measured the distance from each flag to the location of the focal deer prior to flight using a Nikon Prostaff 3i laser rangefinder (Nikon, Inc., Melville, NY). If the approacher, on an angled approach, passed the point tangential to the deer (i.e. began walking away from the deer), the encounter was recorded as a non-flight.

